# An Efficient Lasso Framework for Admixture-Aware Polygenic Scores

**DOI:** 10.1101/2025.08.26.671106

**Authors:** Franklin Ockerman, Brian Chen, Quan Sun, Elena V. Kharitonova, Walter Chen, Laura Y. Zhou, Ruth J.F. Loos, Charles Kooperberg, Ulrike Peters, Jeffrey Haessler, Alexander Reiner, Su Yon Jung, JoAnn E. Manson, Rami Nassir, Kari E. North, Steven Buyske, Christopher A. Haiman, David V. Conti, Lynne R. Wilkens, Ethan M. Lange, Nancy J. Cox, Hongyuan Cao, Laura M. Raffield, Yun Li, Ran Tao

## Abstract

Polygenic scores (PGS) have promising clinical applications for risk stratification, disease screening, and personalized medicine. However, most PGS are trained on predominantly European ancestry cohorts and have limited portability to external populations. While cross-population PGS methods have demonstrated greater generalizability than single-ancestry PGS, they fail to properly account for individuals with recent admixture between continental ancestry groups. GAUDI is a recently proposed PGS method which overcomes this gap by leveraging local ancestry to estimate ancestry-specific effects, penalizing but allowing ancestry-differential effects. However, the modified fused LASSO approach used by GAUDI is computationally expensive and does not readily accommodate more than two-way admixture. To address these limitations, we introduce HAUDI, an efficient LASSO framework for admixed PGS construction. HAUDI re-parameterizes the GAUDI model as a standard LASSO problem, allowing for extension to multi-way admixture settings and far superior computational speed than GAUDI. In extensive simulations, HAUDI compares favorably to GAUDI while dramatically reducing computation time. In real data applications, HAUDI uniformly out-performs GAUDI across 18 clinical phenotypes, including total triglycerides (TG), C-reactive protein (CRP), and mean corpuscular hemoglobin concentration (MCHC), and shows substantial benefits over an ancestry-agnostic PGS for white blood cell count (WBC) and chronic kidney disease (CKD).

## Introduction

Polygenic scores (PGS), which aggregate genetic effects across the genome to generate phenotypic predictions, have demonstrated predictive value for many traits of interest. PGS has potential as a clinical approach to augment existing screening procedures using disease risk stratification^1–4^. However, important barriers to clinical implementation remain, including limited utility for some traits, difficulty in interpretation, and patient concerns over use of genetic data^5–7^. One problem that has received significant attention is the issue of PGS portability, where PGS constructed in one population have attenuated performance when applied to more genetically distant populations^8^. Recent work has demonstrated decreased PGS performance with increasing genetic distance from the training cohort, even within populations typically considered “homogeneous”^9^. Given existing disparities in numbers of genetic samples between genetic ancestry groups, applying PGS without addressing portability has the potential to exacerbate existing health inequities^8^.

The sources of the PGS portability impediment are a subject of considerable debate, but common explanations include gene-gene and gene-environment interactions, population-specific linkage disequilibrium (LD), and minor allele frequency (MAF) differences, including population-specific causal variants^10–12^. In the case of gene-gene and gene-environment interactions, the association between a variant and a phenotype may differ between populations because of differing genetic backgrounds or environments. Similarly, population-specific patterns of LD can cause a tagging variant to correlate highly with a causal variant in one population but have little correlation in another. This yields differing average effects of the tagging variant in each population. In the case of population-specific variants, differences in minor allele frequency between populations can cause variants which are an important source of phenotypic variance in one population to have little explanatory power in another population. In the first two cases (gene-gene/environment interactions and LD differences), the appropriate effect estimate to apply for an individual for PGS depends on their local ancestry, global ancestry, and environment. In the case of MAF differences, the average variant effect is identical between populations, and the effect may be estimated with higher precision from a population where the risk-increasing allele is present in greater frequency.

Recent efforts have focused on cross-population PGS, which typically function by estimating population-specific PGS while borrowing information from external genome-wide association studies (GWAS)^10–13^. In the single-ancestry setting, penalized regression and Bayesian methods have emerged as the dominant approaches^14^. For example, Bayesian methods like LDpred2^15^ and PRS-CS^16^ have enjoyed widespread adoption, while penalized regression methods like Lassosum^17^ have demonstrated scalability and flexibility for diverse genetic architecture. Both paradigms have also been extended to the cross-ancestry setting as well. For example, PRS-CSx is a popular Bayesian method which uses a shared continuous shrinkage prior to integrate GWAS summary statistics from multiple populations and develop population-specific PGS^13^. Likewise, PROSPER is an ensemble method which combines penalized regression models from single-ancestry PRS to generate cross-ancestry PGS^18^. Since genetic architectures are broadly similar across populations^19^, cross-population PGS can increase predictive accuracy for many traits. However, where population-specific variant effects occur, cross-population methods are inappropriate for individuals with recent admixture. For these individuals, whose genetic ancestry varies across the genome, no population-specific PGS will adequately capture the genetic contribution to their phenotypes. For this reason, admixed individuals may be particularly vulnerable to PGS portability issues.

Several recent methods have been proposed to address the limitations of single-ancestry and cross-ancestry PGS in admixed individuals. Marnetto et. al 2020 introduced partial polygenic scores (pPRS), a summary-statistic based PGS method explicitly designed for admixed individuals^20^. PPRS works by segmenting a haplotype into regions based on inferred local ancestry, then applying the appropriate ancestry-specific effect for each variant using external GWAS summary statistics. However, PPRS cannot leverage the unique LD patterns by population which may modify the average allelic effect for admixed individuals. Similarly, variant effects are largely shared between continental ancestry groups, and PPRS does not take advantage of this genetic correlation to improve effect estimation. In the Bayesian paradigm, Zhou et. al recently proposed SDPR_admix^21^, which leverages local ancestry inference in individual-level data to calculate PGS in individuals with 2-way admixture. SDPR_admix characterizes the joint distribution of ancestry-specific effect sizes as a mixture of distributions. Latent components of this distribution represent the cases where both effects are zero, effects are ancestry-enriched (only one is non-zero), or effects are shared with some genetic correlation. Under various simulated and real-data scenarios, SDPR_admix showed superior performance to existing admixed PGS methods, and it was able to scale to biobank-scale datasets.

To address the limitations of existing PGS methods for individuals from underrepresented groups, we recently proposed GAUDI (Genetic Ancestry Utilization in polygenic risk scores for admixed Individuals), a novel method designed for individuals with recent admixture between continental ancestry groups^22^. GAUDI is a penalized regression method which explicitly models ancestry-specific effects and uses a fused LASSO approach to shrink these effects together while encouraging sparsity. GAUDI then estimates individual PGS by applying these effects according to an individual’s mosaic of continental ancestry background across their genome. GAUDI has the advantage of accounting for local patterns of genetic admixture within individuals, as opposed to methods which can only account for global ancestry. In simulated scenarios with ancestry-specific differential effects, GAUDI outperformed existing PGS methods such as pPRS^20^, PRS-CSx^13^, and PRSice^23^. Likewise, GAUDI demonstrated superior performance for traits with known ancestry-specific genetic architectures, such as white blood cell count, in individuals with European (EUR) and African (AFR) admixture. However, the fused LASSO algorithm implemented in GAUDI is computationally demanding and not scalable to large-scale GWAS training datasets. For traits with high polygenicity, i.e., many causal variants, it is infeasible to model a sufficiently large number of variants. Likewise, extension to multiway admixture scenarios, which requires estimating additional parameters for each variant, quickly becomes infeasible under the fused LASSO framework.

Building on the penalization scheme developed in GAUDI, we now introduce HAUDI (Higher-dimensional GAUDI), a computationally efficient PGS method designed for individuals with recent genetic admixture. Like GAUDI, HAUDI is a penalization method that balances sparsity in genetic effects and fusion between ancestry-specific effects. However, HAUDI is reparametrized as a standard LASSO problem, allowing for dramatic improvements in computational performance and extension to multi-way admixture settings. HAUDI is implemented with the commonly-used bigstatsr R package and uses its highly efficient LASSO solver, which accommodates datasets that are too large to fit in memory^24^. In this study, we evaluate HAUDI and compare its performance to GAUDI and the standard LASSO PGS. We also provide a comparison with SDPR_admix, which represents the current state of the art for Bayesian admixed PGS. Using thorough simulation studies and real data analysis on an array of clinical traits, we demonstrate that HAUDI maintains high predictive value for admixed participants while requiring far fewer computational resources than GAUDI.

## Results

### Method Overview

HAUDI is a PGS method designed specifically to accommodate admixed samples. Using inferred local ancestry from individual level data, HAUDI calculates PGS with a penalized regression approach. It reduces to a LASSO model, with coefficients corresponding to 1) the total effect of allele dosage on the phenotype and 2) differences in ancestry-specific allele dosage on the phenotype. With this parameterization, HAUDI applies the standard LASSO penalty to coefficients of the first case and a LASSO penalty with an additional penalty to coefficients of the second case. In this way, HAUDI enforces sparsity in ancestry-specific allelic effect estimates while penalizing differences in ancestry-specific effects. In the standard workflow, reflected in Fig 1), HAUDI begins with estimation of local ancestry, leveraging target genotype data and a panel of external genotypes corresponding to reference ancestries. After estimating local ancestry, HAUDI proceeds by generating a binary file with ancestry-specific allele dosages for variants of interest (either genome-wide or obtained from an external GWAS). This data is accessed using the bigstatsr R package^24^, which uses memory-mapping to accomplish matrix operations on files too large to fit in memory. Finally, HAUDI calculates polygenic scores using the described penalized regression approach.

**Fig 1:**
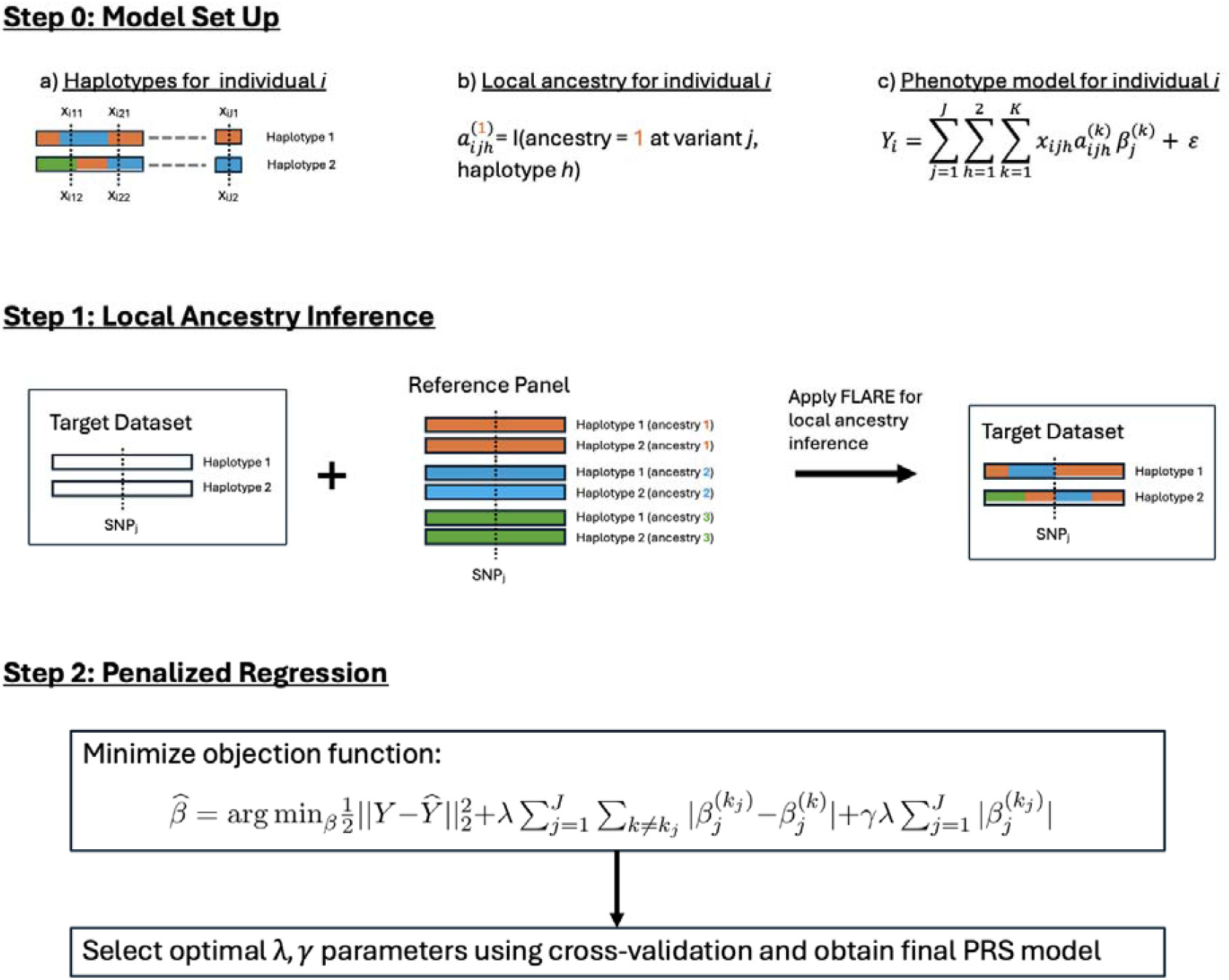
Schematic overview of a HAUDI. In step 0, we define the phenotype model for an individual under the HAUDI framework. HAUDI requires individual-level phased genotype data (a). We assume an individual haplotype is a composition of segments inherited from two or more source populations, and in b), we define an indicator for local ancestry at each variant and haplotype. An individual’s genetic contribution to their phenotype c) is the sum of ancestry-specific effects across the genome. In step 1, we estimate local ancestry in a target dataset, using a reference panel of haplotypes from each source population. In step 2, we minimize HAUDI’s objective function and obtain a set of ancestry-specific effect estimates. We obtain a final PGS by applying these effects to the phenotype model defined in step 0.

### PGS performance in simulated data

We evaluated HAUDI in simulations under a variety of genetic architectures to understand how it compares to GAUDI and the ancestry-agnostic LASSO model (Methods). We observed that all models performed similarly with mean relative difference in R^2^ <3% (Figure 2) when the genetic correlation between CEU and YRI variant effects was 1. In this setting, there is no benefit to modeling local ancestry, so we expect the LASSO model to be non-inferior to the admixed PGS methods (HAUDI, GAUDI, and SDPR_admix). LASSO had small relative improvements in *R*^2^ over GAUDI and SDPR_admix – 1.27% and 2.89%, respectively. LASSO showed only a 0.12% relative improvement over HAUDI in this setting, indicating that HAUDI may be more robust to settings where there is no difference in ancestry-specific effects. In this setting, all models were able to explain a very large proportion of the heritability (>84% for h^2^ = 0.2, >96% for h^2^ = 0.6).

**Fig 2:**
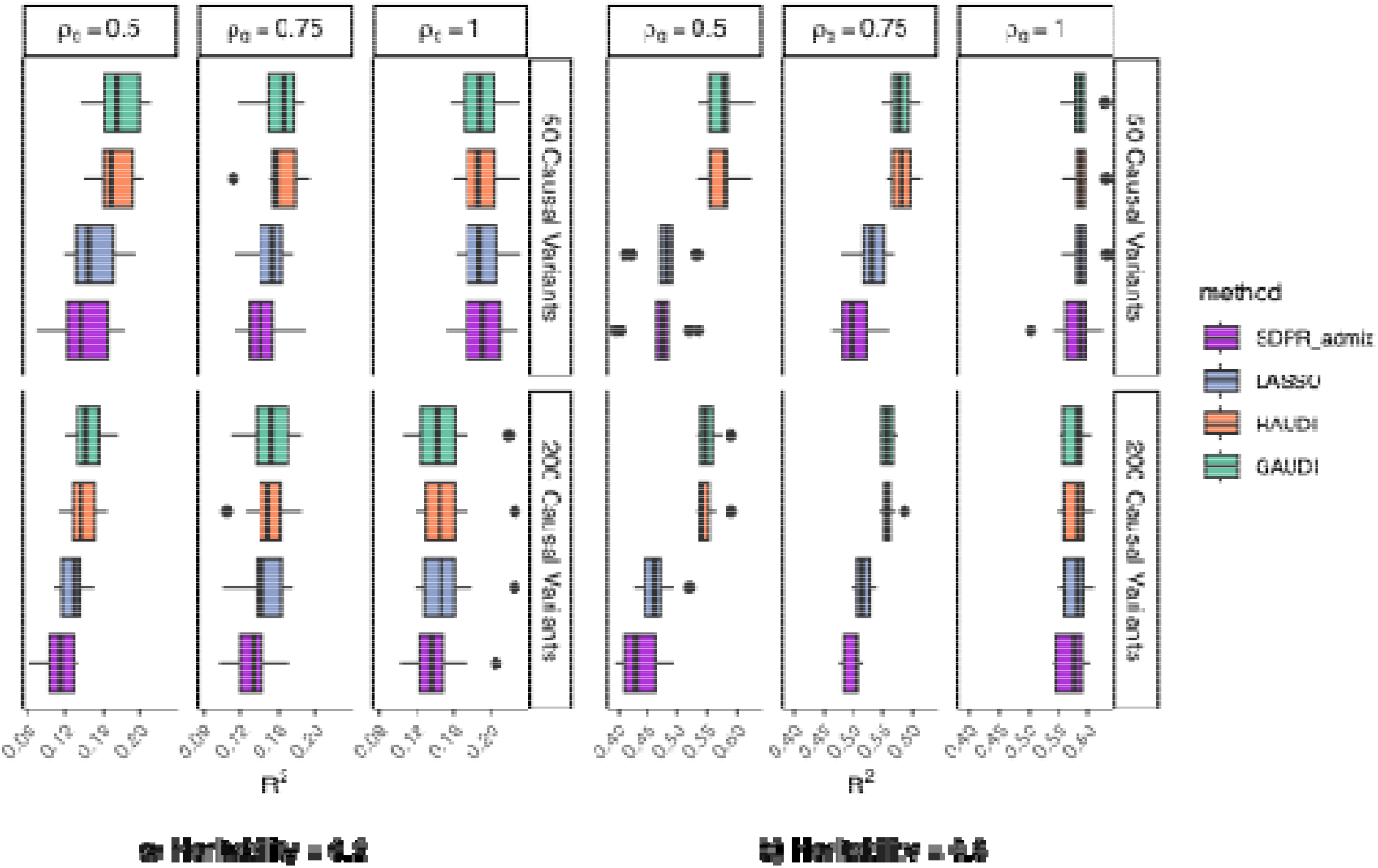
Comparison of PGS methods under various simulation settings. Box-plots correspond to R^2^ value across 10 test sets. Simulation settings include genetic correlation (ρ_g_) between CEU/YRI-specific effects, heritability, and number of true causal variants. All models were fit on 1000 total variants, including causal variants.

When the genetic correlation was 0.5, there were substantial benefits to modeling local ancestry. In this case, HAUDI showed a 15.8% improvement in R^2^ over the ancestry-agnostic LASSO model, while GAUDI showed a 17.9% improvement. In contrast, SDPR_admix performed worse than the LASSO model in this setting, and HAUDI and GAUDI had relative improvements over SDPR_admix of 24.3% and 36.6%, respectively. Both HAUDI and GAUDI maintained similar R^2^ to the corresponding simulations where genetic correlation was 1. However, the mean R^2^ for the LASSO model dropped by 18% when genetic correlation was reduced from 1 to 0.5, indicating that the LASSO model was unable to maintain high accuracy in the presence of ancestry-specific effects. For genetic correlation of 0.75, there were still benefits to modeling local ancestry. Here, HAUDI and GAUDI showed 7.1% and 7.6% improvements, respectively, over the LASSO model.

We hypothesize that the decrease in SDPR_admix’s performance for settings with genetic correlation < 1 is a result of the model failing to distinguish settings with ancestry-specific effects (i.e. only one ancestry has a non-zero SNP effect) from settings where both ancestries have non-zero effects. In settings with genetic correlation equal to 0.5, the average (across simulations) SDPR_admix posterior mode estimate for the proportion of variants with ancestry-specific effects was 0.22. However, the true proportion is 0, since we only perform simulations where both ancestries have the same causal variant.

### Contrasting HAUDI and GAUDI performance in simulated data

In general, HAUDI and GAUDI performed very similarly. Across all simulations, the mean relative improvement in R^2^ of GAUDI over HAUDI was less than 0.5%. We observed the largest difference between the two methods for the setting with h^2^ = 0.2, 200 causal variants, and a genetic correlation of 0.5. Here, HAUDI showed a 4.7% relative reduction in R^2^ from GAUDI. The second largest difference was observed for the setting with h^2^ = 0.2, 200 causal variants, and a genetic correlation of 1. Here, HAUDI showed a 2.9% improvement in R^2^ over GAUDI. Despite the similar performance between HAUDI and GAUDI, the GAUDI effect estimates were noticeably sparser than HAUDI’s (Supplementary Table 3). This was likely due to the Cross-Model Selection and Averaging (CMSA) scheme used in HAUDI’s LASSO solver, in which HAUDI effect estimates were obtained by averaging over inner cross-validation folds rather than re-fitting on the full training set^20^. For simulations with 50 causal variants (out of 1000), 59.5% of HAUDI effect estimates were equal to 0 (on average), compared with 84.7% for GAUDI. For simulations with 200 causal variants, these figures were 32.6% and 65.2%, respectively. Overall, HAUDI showed a major improvement in computational efficiency compared to GAUDI. Across all simulations, the median run time was 25 seconds for HAUDI and 480 minutes for GAUDI – a >99% reduction in run time (Supplementary Figure S3).

### Ancestry-specific estimates in simulated data

We compared the true CEU-specific effects versus estimated CEU-specific effects in all models to evaluate bias in effect estimates (Figure 3). We observed similar performance across models when the genetic correlation between CEU and YRI variant effects was set to 1. In this case, where true CEU and YRI-specific effects were equal, the correlation between true and estimated CEU-specific effects was greater than 0.92 for all methods. When the genetic correlation was 0.5, HAUDI and GAUDI maintained high correlation (0.82 and 0.84 respectively) between true and estimated CEU-specific effects, but the correlation for the LASSO and SDPR_admix models dropped to 0.61 and 0.58, respectively. In turn, the scatter plot for the LASSO and SDPR_admix models show larger deviation between true and estimated effects than scatter plots for HAUDI and GAUDI. These results are expected for LASSO, since the LASSO model is unable to distinguish ancestry-specific effects. For YRI-specific effects (provided in Supplementary Figure S2), this effect was somewhat diminished, and the LASSO model retained a higher correlation of 0.81. It is likely that the LASSO model was able to capture YRI-specific effects better than CEU-specific effects because of the higher overall representation of YRI haplotypes in the simulated data. Since the ancestry-agnostic LASSO model forces ancestry-specific effects to be equal, these estimates are dominated by the YRI-specific effects.

**Fig 3:**
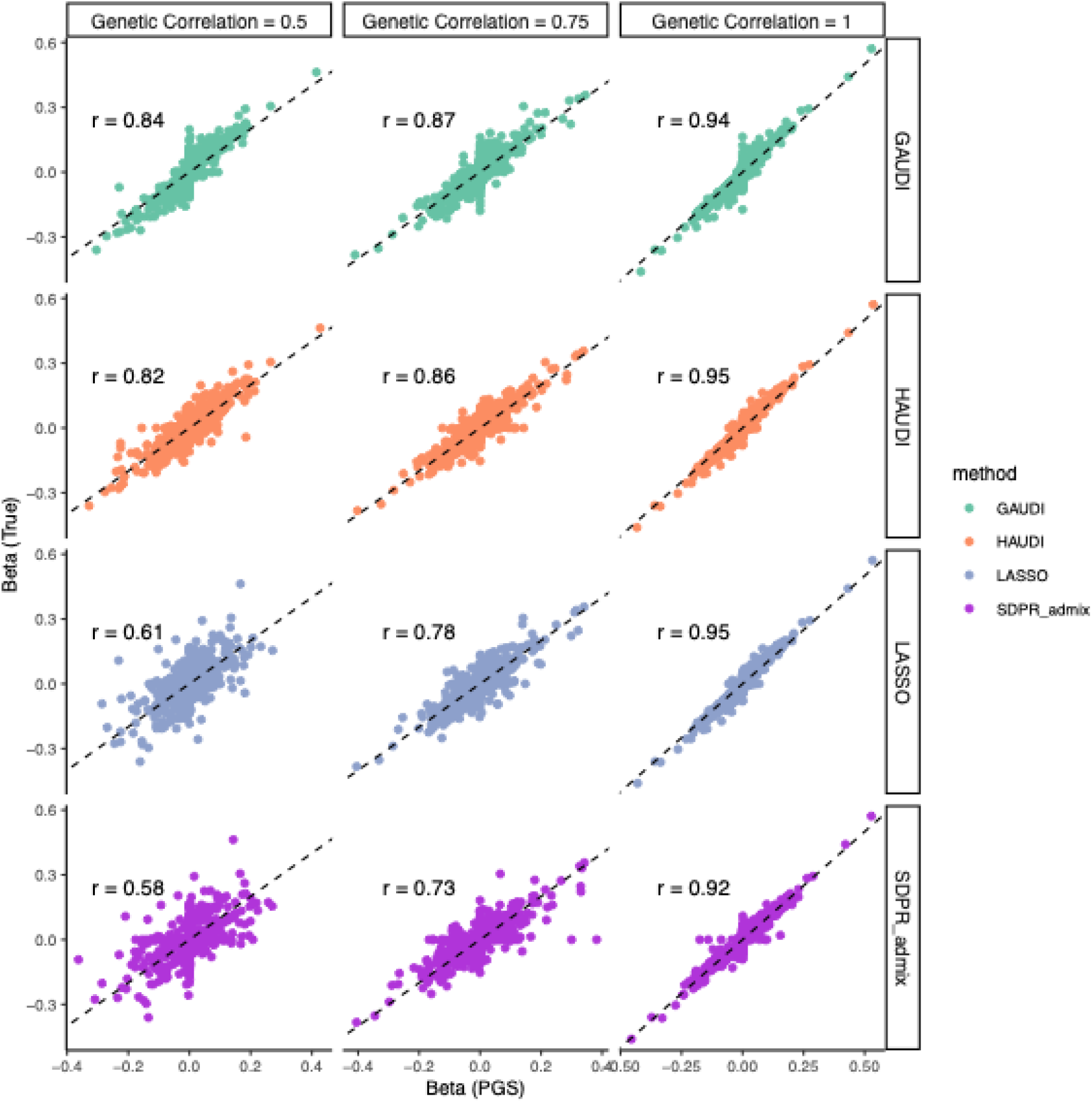
Comparisons between estimated and true CEU-specific effects. Plotted in each panel are the estimated (x axis) and true (y axis) variant effects, stratified by genetic correlation and PGS method. Only the first repetition is plotted for illustration, however the correlation between the true and estimated effects across all simulations (r) is annotated for each panel. This figure is restricted to CEU-specific effects. For LASSO, the x-axis shows estimated ancestry-agnostic effects.

### Analyses in PAGE African American Participants

To assess HAUDI’s performance on real data, we evaluated PGS models on 19 clinical traits, using self-reported African American and Hispanic/Latino individuals from the PAGE Study. No African-specific Pan-UK Biobank GWAS summary statistics were available for QRS interval, so this phenotype was omitted from the analysis of PAGE African American participant data.

Results for the real data analysis on PAGE African American subjects are reported in Figure 4, where we fit penalized regression models (HAUDI, GAUDI, LASSO) using the top 500 pruned variants from the UK Biobank GWAS. This allows for a fair comparison to GAUDI, which is too computationally intensive to fit on tens of thousands of variants. For SDPR_admix, we fit models using the top 50,000 variants. A comparison of run times (aggregated across chromosomes and tuning parameters) is provided in Figure 5. For SDPR_admix models fit with 50,000 variants, the mean (median) run time was 223.1 (115.0) minutes. For HAUDI, the mean (median) run time for models fit with 50,000 variants was 16.4 (11.2) minutes, representing an average reduction of 86% in wall clock time. The SDPR_admix average is heavily skewed by large run times for PR interval (mean of 1941.5 minutes). However, even omitting this phenotype, the average reduction in wall clock time for HAUDI vs. SDPR_admix was 85.2%.

**Fig 4:**
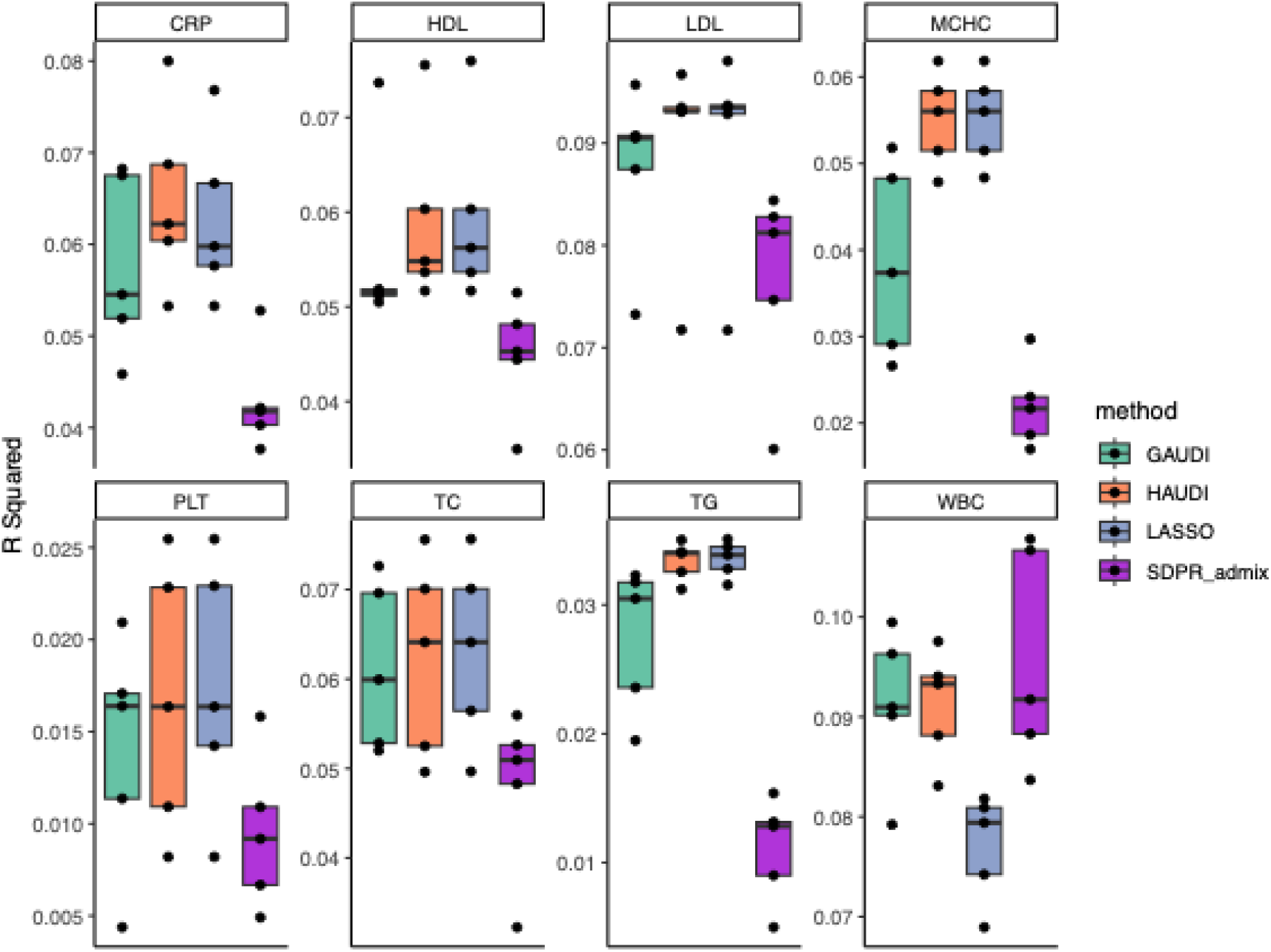
Comparison of test set R^2^ on PAGE African American data. Penalized regression models (GAUDI, HAUDI, LASSO) were fit using the top 500 pruned variants by p-value in UKB GWAS data. SDPR_admix was fit using the top 50,000 pruned variants. Phenotypes are restricted to those with N > 5000 and mean R^2^ > 0.01 in at least one model. Per-phenotype boxplots correspond to the R^2^ values in each test set fold.

**Fig 5:**
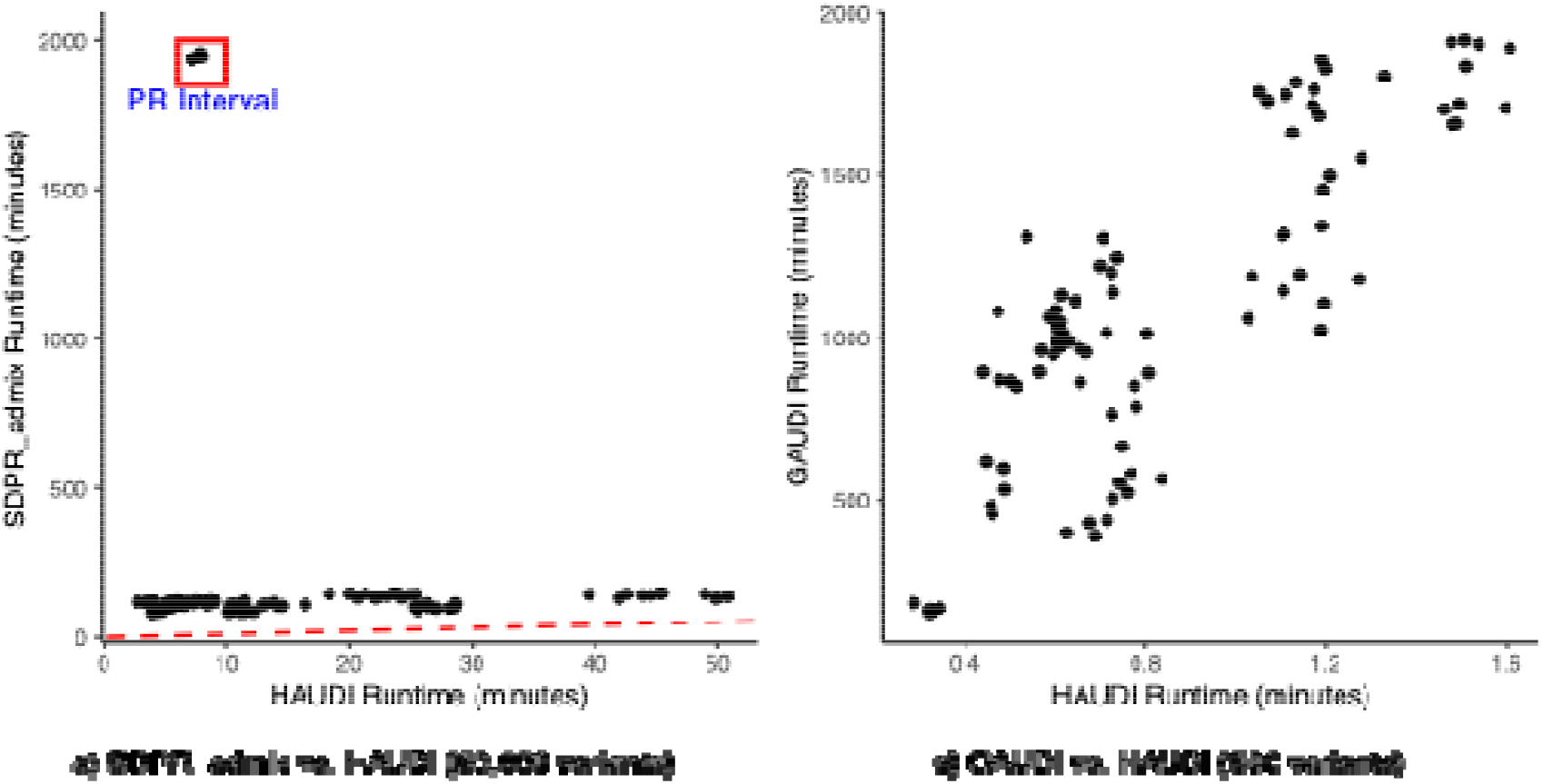
Comparison of PGS run time on data from PAGE African American participants. We report run times across phenotypes and testing folds for the three admixed PGS models. To compare SDPR_admix and HAUDI (a), we restrict to models fit on 50,000 variants. To compare GAUDI and HAUDI, we restrict to models with 500 variants.

For GAUDI models fit with 500 variants, the mean (median) run time was 1101.2 (1056.0) minutes, vs a mean of 0.86 minutes and a median of 0.73 minutes for HAUDI. This represents a > 99.9% improvement in run time.

Among the 8 phenotypes where at least one method reached mean *R*^2^ > 0.01, HAUDI performed best, with mean relative improvements in *R*^2^ over GAUDI (16.3%), LASSO (1.7%) and SDPR_admix (80.4%). For mean corporeal hemoglobin concentration (MCHC), platelet count (PLT), and triglycerides (TG), we observe notably diminished performance in GAUDI, with mean relative decreases in *R*^2^ compared to HAUDI of 29.9%, 16.9%, and 17.5%, respectively. For SDPR_admix, the same three traits showed the largest difference in performance compared to HAUDI, with relative decreases in *R*^2^ of 159% (MCHC), 100.3% (PLT), and 254% (TG).

HAUDI showed similar performance to LASSO for 7 out of 8 traits (mean relative difference in *R*^2^ <5%), indicating that modeling local ancestry did not uniformly improve predictive power. However. for white blood cell count (WBC), all three admixed PGS methods out-performed LASSO. SDPR_admix performed best for this trait, with a mean test-set *R*^2^ of 9.6% and relative improvement over LASSO of 24.0%. Similarly, HAUDI (mean *R*^2^ = 9.1%) had a 18.5% improvement over LASSO, and GAUDI (mean *R*^2^ = 9.1%) had an 18.6% improvement over LASSO. For the other 7 phenotypes, where modeling local ancestry did not significantly improve predictive power, HAUDI maintained a large advantage over GAUDI (18.5% improvement) and SDPR_admix (92.5% improvement). This indicates that HAUDI performs similarly to GAUDI and SDPR_admix in settings where ancestry-specific effect differences influence phenotypic variance, and it substantially improves over them in the null case where modeling local ancestry does not increase in-sample *R*^2^.

### Analyses in PAGE Hispanic/Latino Participants

For PAGE Hispanic/Latino participants, we did not fit models for GAUDI or SDPR_admix, as they do not accommodate more than 2-way admixture scenarios. Instead, we compare HAUDI vs the ancestry-agnostic LASSO PGS. Without the computational limits imposed by GAUDI, we chose to fit models using the top 500, 10000, or 50000 pruned variants from the UK Biobank GWAS, selecting the best model per training fold with internal cross-validation. Box-plots for PGS *R*^2^ in held-out test sets are provided in Figure 6. As in the PAGE African American dataset, we restrict results to phenotypes with mean *R*^2^ > 0.01 in at least one method. Across the 13 remaining phenotypes, there was a small improvement in *R*^2^ for HAUDI vs. LASSO (2.7%). However, for white blood cell count and chronic kidney disease (CKD), there were large benefits, with corresponding improvements of 31.9% and 7.6%. For PR interval, HAUDI performed slightly worse than LASSO, with a mean relative decrease in R2 of −2.2%.

**Fig 6:**
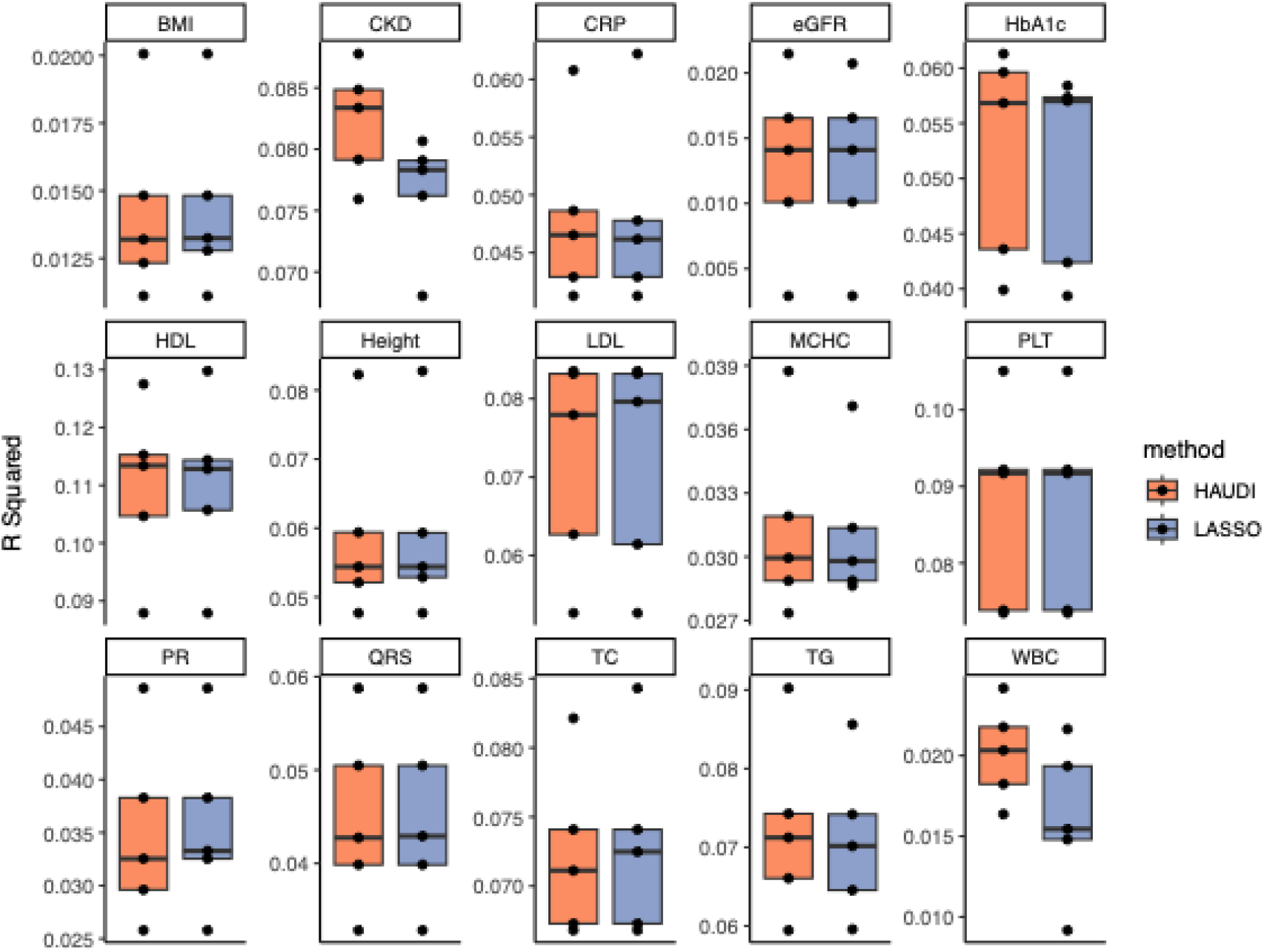
Comparison of GAUDI and LASSO test set R^2^ on PAGE Hispanic/Latino data. All models were fit using the top 500, 10000, or 50000 variants (by p-value in UKB GWAS data). Phenotypes are restricted to those with N > 5000 and mean R^2^ > 0.01 in at least one model. Per-phenotype boxplots correspond to the R^2^ values in each test set fold.

### Ancestry-Specific Effects in PAGE Data

It is notable that modeling local ancestry improved R^2^ for white blood cell count over the ancestry-agnostic LASSO model in both the African American and Hispanic/Latino datasets. Previous studies have observed that benign neutropenia, characterized by low white blood cell counts without apparent adverse effects, is common in individuals with African ancestry^25,26^. It has been noted that individuals with what has been previously referred to as “benign ethnic neutropenia” (more recent papers propose the term “Duffy-null associated neutrophil count”, as individuals with multiple self-identified ethnicities may be impacted) are at risk for inappropriate clinical care, an effect which may exacerbate existing health inequities^27,28^. Multiple genetic studies have identified the Duffy null phenotype as an important predictor of benign neutropenia. This phenotype is caused by the Duffy null variant (rs2814778) at the promoter for the *ACKR1* gene, which is common in African (>95%) but uncommon in European(<5%) reference panels^25,27,29^. For African-American PAGE participants, three out of four variants with the largest differences in ancestry-specific effect sizes (rs863029, rs1633247, rs1772444), were located within 200kb of the Duffy null variant. Likewise for the Hispanic/Latino PAGE participants, the variant with the largest difference in ancestry-specific effect sizes (rs12075) was located less than 1kb from the Duffy null variant. This is likely a result of these variants tagging the Duffy null variant in African haplotypes but not in European haplotypes. To illustrate the pattern of ancestry-specific effects for white blood cell count, we report scatter plots in Figure 7 (African-American participants) and Figure 8 (Hispanic-Latino participants).

**Fig 7:**
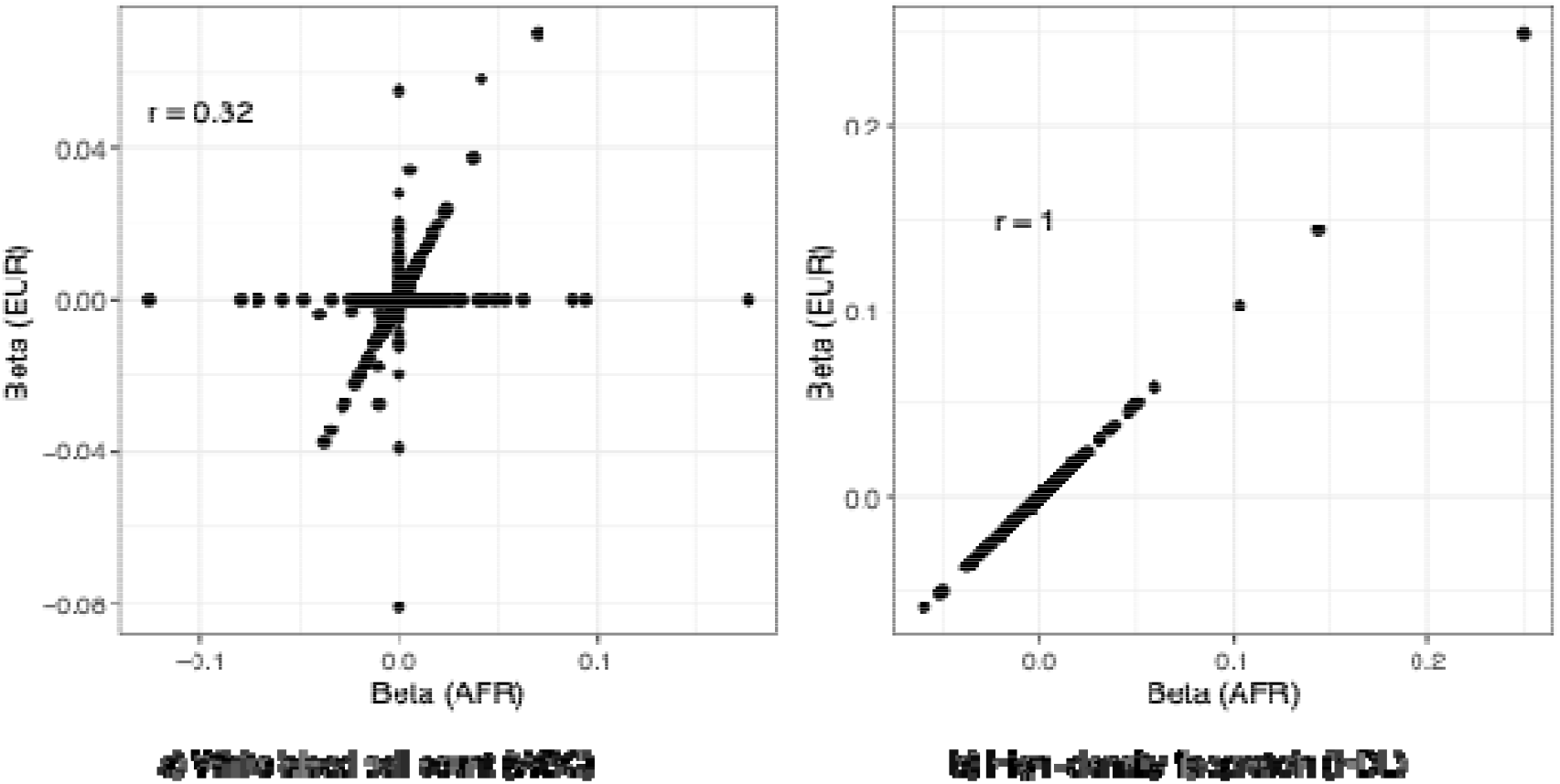
Comparison of HAUDI ancestry-specific effect estimates (African-American participants). Effect estimates were obtained using the optimal set of tuning parameters (selected by cross-validation) in the first training fold. HDL (b) was chosen as a comparator for WBC (a) to demonstrate that HAUDI can also capture genetic architectures with similar effects across ancestries. Correlation between ancestry-specific effects (r) is given in each panel.

**Fig 8:**
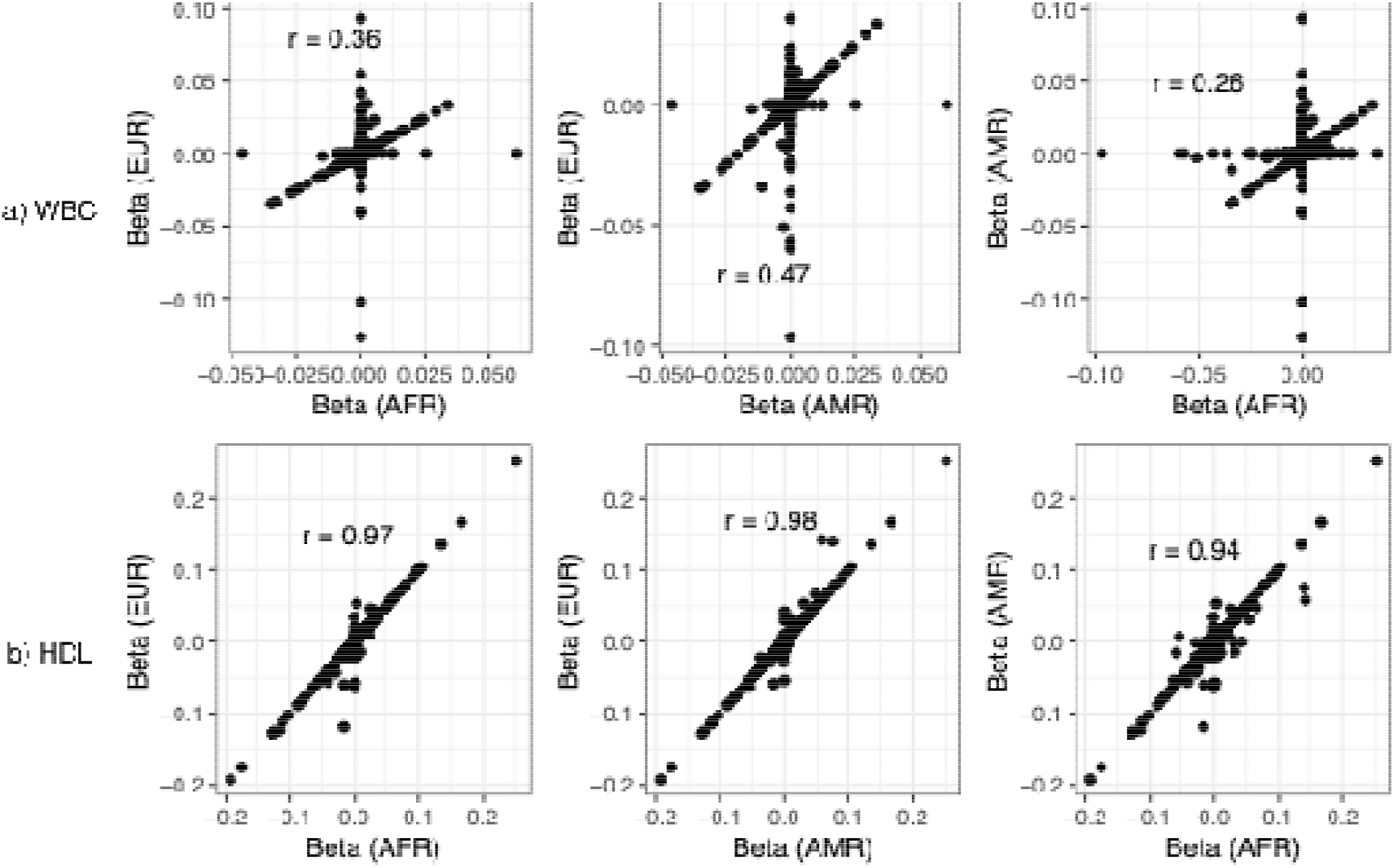
Comparison of HAUDI ancestry-specific effect estimates (Hispanic/Latino samples) for White Blood Cell Count (WBC). Effect estimates were obtained using the optimal set of tuning parameters (selected by cross-validation) in the first training fold. HDL (b) was chosen as a comparator for WBC (a) to demonstrate that HAUDI can also capture genetic architectures with similar effects across ancestries. Correlation between ancestry-specific effects (r) is given in each panel.

## Discussion

Polygenic scoring has emerged as an exciting methodological approach for phenotype prediction in clinical settings. For the promises of PGS to be realized, it is critical that investigators and clinicians adopt methods which are effective for all individuals, including those with recent continental admixture. In this work, we proposed and evaluated HAUDI, an upgrade of our previous method GAUDI, which is designed to incorporate inferred local ancestry in the estimation of ancestry-specific variant effects. In traits where LD patterns or population-specific variants limit PGS generalizability, HAUDI can improve predictive performance for admixed individuals by allowing for ancestry-specific effect estimates. For traits with consistent genetic architecture across ancestry groups, HAUDI flexibly penalizes differences in ancestry-specific effects. In addition, HAUDI is highly computationally efficient and accommodates multi-way admixture settings. While no “one-size fits all” approach exists for PGS, HAUDI is a promising approach for improving PGS performance in individuals of diverse ancestry.

In simulated data, HAUDI out-performed GAUDI in the null setting where no ancestry-specific effects exist and genetic architecture is identical across ancestry groups. HAUDI yielded effect estimates and predictive performance consistent with an ancestry-agnostic LASSO comparator, indicating that HAUDI is robust to settings where local ancestry is uninformative. When we introduced population-specific variant effects, HAUDI was able to capture these differences and showed substantial improvements over the ancestry-agnostic comparator. In these cases, HAUDI performance was consistent with GAUDI while requiring far fewer computational resources. These results were replicated in real data applications, where HAUDI out-performed GAUDI for a variety of clinical phenotypes and meaningfully improved over the ancestry-agnostic comparator for white blood cell count and residualized chronic kidney disease. For white blood cell count, previous studies have implicated the Duffy null variant as a common cause of benign neutropenia in individuals with African ancestry. This variant is highly population-differentiated, being virtually absent in European individuals. These results indicate that HAUDI can effectively capture ancestry-specific variants and effects in admixed individuals

A limitation of this work is that HAUDI requires the use of individual level data. Although this requirement may limit the applicability of HAUDI for some investigators, individual level data is currently necessary to effectively model local ancestry. While methods such as lassosum^17^ use summary statistics to approximate penalized regression, it is currently unclear whether this approach is suitable for application to local ancestry. Future work may focus on whether it is possible to leverage summary-level data for estimating population-specific effects in admixed cohorts. In addition to external GWAS summary statistics, we also hope to address variant function annotations, another important source of external information, in future work. Methods such as SBayesRC^30^ have demonstrated that polygenic prediction can be improved by prioritizing functional variants, which are more likely to be causal. A final limitation is the lack of explicit guidelines for prioritizing variants to include in admixed PGS models. In this work, we performed pruning and thresholding using external GWAS corresponding to the majority ancestry in each dataset. However, alternative methods may include performing GWAS in an admixed cohort of similar composition or meta-analysis of external GWAS from multiple populations. Given the computational efficiency of HAUDI, it may be reasonable in many cases to perform PGS using a large set of common variants without additional pruning and thresholding. The impact of these choices on PGS accuracy in the admixed population setting is unclear, and future work should focus on developing a principled approach to this problem.

HAUDI also relies on accurate inference of local ancestry. Our pipeline assumes input from the FLARE program^31^, which is highly performant and purports high accuracy across a range of admixture histories and genetic architectures^32^. The impact of inferred ancestry accuracy on HAUDI results has not been evaluated in this paper. However, HAUDI is ultimately agnostic to the choice of local ancestry estimation. Alternative methods such as RFMix^33^ may be used, although an ad-hoc approach may be required to convert window-based local ancestry files into the local ancestry-formatted VCF input required by HAUDI.

In summary, we have presented a novel method for computationally efficient polygenic scoring which explicitly accounts for local ancestry in admixed individuals. Using simulated and real datasets, we demonstrated that our method can capture population-specific effects in traits with population-specific genetic architectures while maintaining high predictive value in settings where genetic architectures are shared between ancestries. We believe this work represents an important step towards polygenic scores with clinical utility for all individuals.

## Methods

### The GAUDI Model

The GAUDI model can be summarized as follows: for a set of admixed individuals 1,…,*n* with ancestry from populations 1,…*K* let *x*_*ij*1_ be the allelic value for individual *i* at variant *j* on haplotype 1 (with the same notation for haplotype 2), j=1,2,…,J. Similarly, let a_ij1_ be the local ancestry of individual *i* at variant *j* on haplotype 1. Now letting 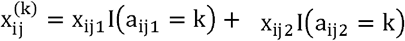,we may consider 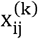 the ancestry-specific genotype dosage for ancestry *k*. Introducing matrix notation, we define the phenotype *Y*, ancestry-specific genotypes *G*, and variant effects β as:

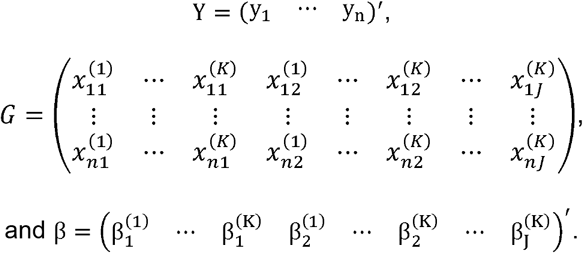

For the purposes of illustration, we assume *Y* is continuous, and *Y* and *G* are centered. With this notation, we define the GAUDI phenotype model as:

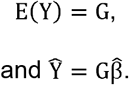

In GAUDI, it is assumed that the admixture occurs between only two ancestral populations, so *K* = 2. With this assumption, we proceed by defining a 3J × 2J penalty matrix *D*:

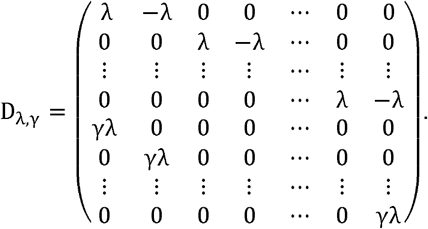

Finally, the GAUDI estimate for β is obtained by solving:

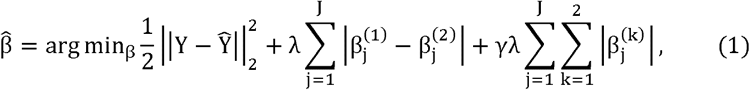

which is equivalent to:

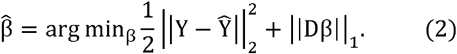

Examining the penalty term in (1), we note that the term 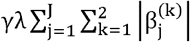 is an *L*_1_ penalty which encourages sparsity in effect estimates. However, the term 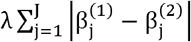 shrinks ancestry-specific effects together, a property known as fusion. Thus, the tuning parameter γ controls the ratio between sparsity and fusion. This penalization scheme is known as a “fused LASSO” model.

### The HAUDI Model

The HAUDI model follows a similar approach to GAUDI to balance fusion and sparsity. For the general case of K ≥2, we denote the most common ancestry at each variant *j* as *k*_*j*_. The most common ancestry at each variant is chosen as the reference ancestry, although this choice is arbitrary. arbitrary. The HAUDI estimate of β is given by:

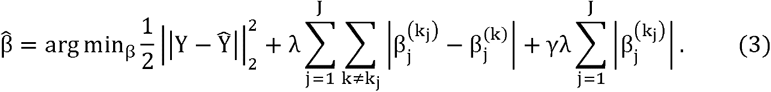

Comparing equations (3) and (1) for the case *K =* 2, the difference is that while GAUDI applies the sparsity penalty γλ to each effect estimate 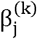, HAUDI only applies this penalty to the effect corresponding to the most common ancestry 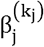 at each locus. Since we no longer directly penalize all ancestry-specific effects, it may appear that HAUDI is less sparse than GAUDI overall. However, sparsity in other ancestry-specific effects is still encouraged due to the penalties on differences in ancestry-specific effects. To understand the practical significance of HAUDI’s modified penalization scheme, we introduce new notation. First, we define:

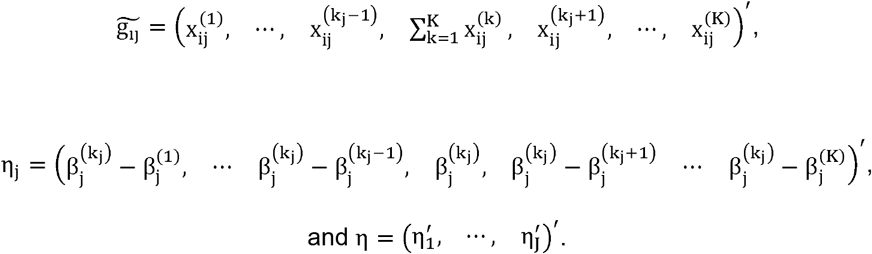

For example, if *K* = 2 and *k*_*j*_ = 1, then 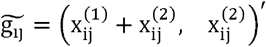 and 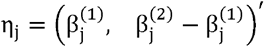. Now, we define the n×(JK) matrix:

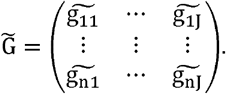

We see that:

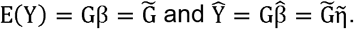

We now define a new JK ×JK penalty matrix, 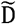:

Let 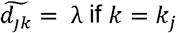 and 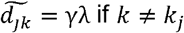. We define the vector of penalties 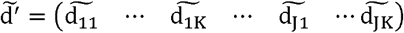 and the associated penalty matrix: 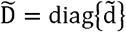.

Finally, we may equivalently express the HAUDI solution in (3) as:

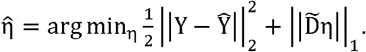

Defining 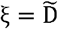 and 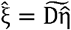, we see that:

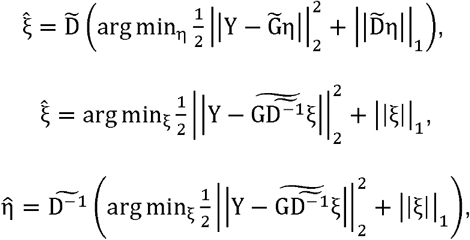

and finally,

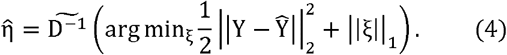

In this way, we see that the re-parameterized model in HAUDI is equivalent to a standard LASSO problem. Because (4) can be solved using standard LASSO solvers, HAUDI is far more computationally efficient than the fused LASSO problem of GAUDI.

### Simulation Study

We simulated recently admixed genotypes using Haptools, implemented with the admix-kit python package^30^. For reference data, we used 1000 Genomes data from 179 Northern European (CEU) and 178 Yoruba African-ancestry (YRI) individuals, subset to chromosome 1 Hapmap3 variants^34^. We simulated 10 generations of admixture with global ancestry proportions of 0.8 and 0.2 for YRI and CEU, respectively. This yielded 10,000 admixed individuals with 86,994 total variants.

After generating admixed genotypes, we simulated phenotypes under a variety of genetic architectures, again using admix-kit^35^. We varied the heritability (20%, 60%), number of causal variants (50, 200), and genetic correlation between the CEU and YRI-specific variant effects (0.5, 0.75, 1). We then split the samples into ten independent testing folds, with corresponding training sets in the left-out samples. This yielded a train/test split of 90%/10%. For parameter tuning in HAUDI, GAUDI, and LASSO, we employed 5-fold cross validation within the training splits.

Due to the higher computational burden of GAUDI, instead of fitting models on all 86,994 variants, we obtained a set containing causal variants (either 50 or 200) and a random sample of non-causal variants (either 950 or 800) for a total of 1000 variants per simulation. For fair comparison across PGS methods, the same variant set for each simulation was used for all PGS models. For each simulated phenotype, we fit models for GAUDI and HAUDI, varying the γ fusion penalty over 0.01, 0.5, 1, 2, and 5. Since genetic correlation is known, we used the true value as input for SDPR_admix models. In addition, we calculated PGS using a standard LASSO model to compare admixed PGS methods with an ancestry-agnostic PGS.

### PAGE Dataset

To test HAUDI on real data, we fit PGS models using data from the Population Architecture using Genomics and Epidemiology (PAGE) Study^36^. Using samples from PAGE participants, we constructed two datasets representing self-identified African American subjects and self-identified Hispanic subjects. The African American data set included 17,300 individuals, consisting of participants from the BioME Biobank Program, Multiethnic Cohort (MEC) Study, and Women’s Health Initiative (WHI). The Hispanic/Latino data set included 11,776 participants from The Hispanic Community Health Study / Study of Latinos (HCHS/SOL). Per-cohort demographic information (age, sex) is provided in Supplementary Table 1. Genotyping for these participants was performed by the PAGE Study using the Multiethnic Genotyping (MEGA) array, an array designed to increase variant coverage across multiple populations^36,37^. Quality control was performed as described previously^36^, such as exclusion for variants with missing call rate >= 2%, high Mendelian error rates, Hardy–Weinberg *p*-value < 1×10−4, large sex differences in heterozygosity or allele frequency, and other criteria. Genotype imputation was performed as described in previous work^38–40^, using the TOPMed freeze 8 reference panel^41^ with minimac4^42^ for imputation and Eagle v2.4^43^ for phasing. To exclude variants of poor imputation quality, we applied a filter based on estimated imputation quality (Rsq) to remove variants with Rsq < 0.6. We additionally removed all variants with minor allele frequency (MAF) < 0.01.

**Table 1:**
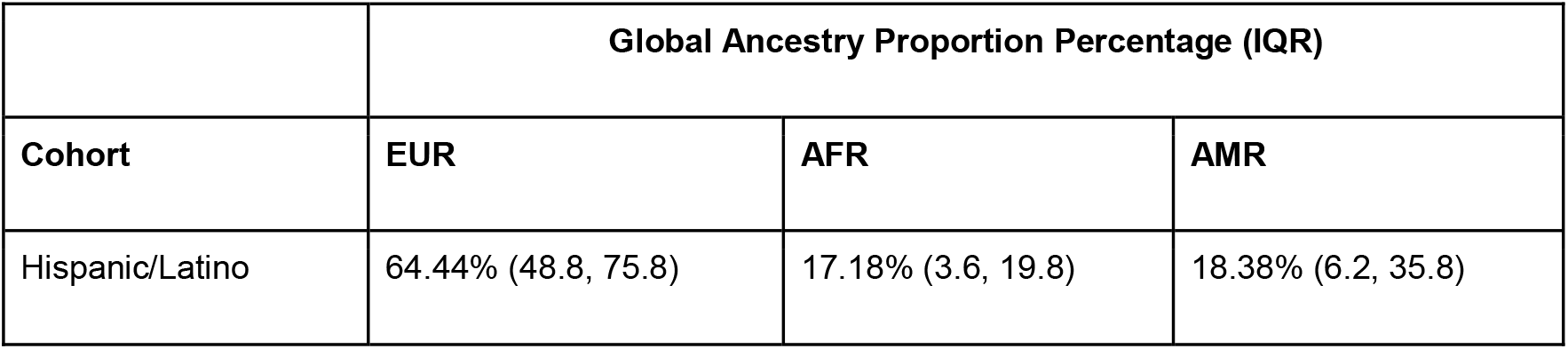
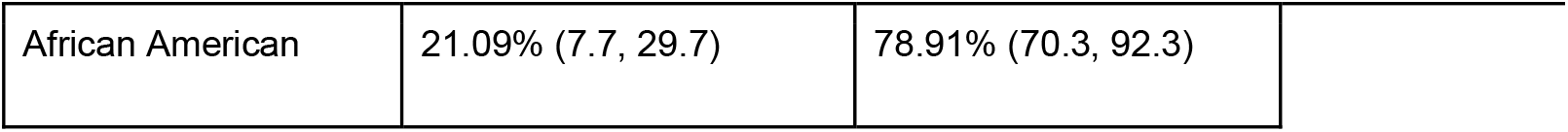
Summary of Global Ancestry Proportions in the PAGE Data. Mean (IQR) global ancestry proportions estimated using FLARE. IQR calculated using individual-level global ancestry proportion estimates.

|  | Global Ancestry Proportion Percentage (IQR) |  |  |
| --- | --- | --- | --- |
| Cohort | EUR | AFR | AMR |
| Hispanic/Latino | 64.44% (48.8, 75.8) | 17.18% (3.6, 19.8) | 18.38% (6.2, 35.8) |
| African American | 21.09% (7.7, 29.7) | 78.91% (70.3, 92.3) |  |

We independently performed local ancestry inference for each data set (African American and Hispanic/Latino) using FLARE^31^. For self-identified African American individuals, we assumed two-way admixture and used a reference panel consisting of 92 individuals of African ancestry and 92 of European ancestry from the 1000 Genomes Project^44^. For self-identified Hispanic/Latino individuals, we assumed three-way admixture and augmented this reference panel with 92 individuals of Native American ancestry from the Human Genome Diversity Project^45^. This reference panel was chosen to balance sample size between each reference population; reference populations were chosen based on prior genetic ancestry analyses of PAGE participants, but we note that two- and three-way admixture models may be a simplification. However, quantitatively smaller local ancestry contributions (for example local ancestry regions similar to Native American reference panels in PAGE African American participants) would be difficult to computationally model. Before running FLARE, we excluded variants with MAF < 0.05 from the reference panels.

### PAGE Real Data Analysis

Based on the availability of external large-sample GWAS summary statistics from the Pan-UK Biobank analyses^46^, we selected 19 clinical phenotypes for analysis. Phenotypes were previously cleaned and harmonized between PAGE studies, and a full description is available in the supplementary materials of Wojcik et. al^36^. Briefly, quantitative phenotypes were adjusted for self-identified race/ethnicity, study/site, and 10 genetic principal components. Additional adjustments were carried out for certain phenotypes, including age, sex, or BMI adjustment. Per-phenotype sample sizes are reported in Supplementary Table 2. Within the two data sets (African American vs. Hispanic/Latino participants), we further adjusted all phenotypes for age, age squared, sex, study, and the first 20 genetic principal components using linear regression for continuous traits and logistic regression for binary phenotypes. After that, we obtained residuals from linear models for continuous traits and deviance residuals from logistic regression models for binary traits. We then obtained the final adjusted phenotypes by performing inverse normal transformation of these residuals.

Due to GAUDI’s higher computational demands, it was necessary to perform p-value clumping and thresholding on variants before fitting PGS models. To do this, we used the Pan-UK Biobank GWAS summary statistics^46^ (see Supplementary Data). We used PLINK^47^ to perform LD clumping within 250kb regions with a *p*-value threshold of 0.05 and an LD threshold of 0.1. For the PAGE African American dataset, we used African-specific UK Biobank GWAS summary statistics to perform clumping and thresholding. For the Hispanic/Latino dataset, we used European-specific UK Biobank GWAS summary statistics. Although we do not assess genetic similarity between PAGE and UKB GWAS samples, this choice reflects the majority global ancestry proportion estimated for each dataset, as reported in Table 1 and Supplementary Figure S1. For fitting PGS, we divided the two datasets into five test sets, with corresponding training sets. This yielded a train/test split of 80%/20%.

PGS were calculated on the training data using HAUDI, GAUDI, SDPR_admix, and an ancestry-agnostic standard LASSO model. For GAUDI and HAUDI, we fit models varying γ over 0.01, 0.5, 1, 2, and 5. For computational reasons, we limited GAUDI models to only the top 500 variants (by *p*-value) for each phenotype. For HAUDI and LASSO, we further defined a grid of models by varying the number of variants between 500, 10000, and 50000 variants. For SDPR_admix, we fit models using the top 50000 variants. Best performing models for each dataset, phenotype, and fold were selected by highest *R*^2^ value in the validation sets. For GAUDI, we used 5-fold internal cross-validation to select the optimal γ parameter for each model. For HAUDI and LASSO, we used 5-fold Cross-Model Selection and Averaging (CMSA) for each model, as defined in the bigstatsr R package^24^.

## Data Availability

Data from the 1000 Genomes Project, used to construct reference panels for local ancestry inference and to simulate admixed genotypes, is publicly available and free to access from the consortium website (https://www.internationalgenome.org/). Data for the Human Genome Diversity Project is publicly available at the same source. GWAS summary statistics for the Pan-UK Biobank, used to select variants for analysis, are publicly available at https://pan.ukbb.broadinstitute.org. Individual-level data, used for constructing and evaluating polygenic scores, can be accessed through dbGaP at https://www.ncbi.nlm.nih.gov/projects/gap/cgi-bin/study.cgi?study_id=phs000356.v2.p1.

## Code Availability

The HAUDI R package is available at https://github.com/frankp-0/HAUDI. All analyses were performed using HAUDI version 0.2.8. A new major version (>1.0.0) is available, with improved computational performance and support for RFMix2, FLARE, and.lanc local ancestry input. Codes used for analyses are available at https://github.com/frankp-0/HAUDI_simulation_analysis and https://github.com/frankp-0/HAUDI_PAGE_analysis.

## Acknowledgments

This study is supported by the National Institutes of Health for the project “Polygenic Risk Methods Development (PRIMED) Consortium,” with grant funding for the EPIC-PRS Study Site (U01HG011720) and CAPE Study Site (U01HG011715). The content is solely the responsibility of the authors and does not necessarily represent the official views of the National Institutes of Health. The PAGE Study is supported by the following NIH grants: U01HG007419, R01HG010297, and R01HL151152. Additional support includes R01AR083790, R01HL146500, and U24AR076730. The authors thank the staff and participants of HCHS/SOL for their important contributions. Investigators website - http://www.cscc.unc.edu/hchs/. The Hispanic Community Health Study/Study of Latinos is a collaborative study supported by contracts from the National Heart, Lung, and Blood Institute (NHLBI) to the University of North Carolina (HHSN268201300001I / N01-HC-65233), University of Miami (HHSN268201300004I / N01-HC-65234), Albert Einstein College of Medicine (HHSN268201300002I / N01-HC-65235), University of Illinois at Chicago (HHSN268201300003I / N01-HC-65236 Northwestern Univ), and San Diego State University (HHSN268201300005I / N01-HC-65237). The following Institutes/Centers/Offices have contributed to the HCHS/SOL through a transfer of funds to the NHLBI: National Institute on Minority Health and Health Disparities, National Institute on Deafness and Other Communication Disorders, National Institute of Dental and Craniofacial Research, National Institute of Diabetes and Digestive and Kidney Diseases, National Institute of Neurological Disorders and Stroke, NIH Institution-Office of Dietary Supplements. We also acknowledge all participants from the WHI program, which is funded by the National Heart, Lung, and Blood Institute, National Institutes of Health, U.S. Department of Health and Human Services through contracts 75N92021D00001, 75N92021D00002, 75N92021D00003, 75N92021D00004, 75N92021D00005.

