## Supplementary Information for "An Efficient Lasso Framework for Admixture-Aware Polygenic Scores"

|  | **WHI (N=6820)** | **BioME(N=5155)** | **MEC (N=5325)** | **HCHS/SOL (N=11776)** |
| --- | --- | --- | --- | --- |
| **Age (Mean/SD)** | 60.00 (6.74) | 52.19 (14.29) | 60.39 (8.58) | 46.09 (13.84) |
| **Sex (%Male)** | 0% | 37.56% | 37.22% | 41.42% |

**Table 1: Summary of Cohort Demographic Information**

| **Phenotype** | **Abbreviation** | **N WHI (N cases)** | **N BioME (N cases)** | **N MEC (N Cases)** | **N HCHS/SOL (N Cases)** |
| --- | --- | --- | --- | --- | --- |
| Height | Height | 6808 | 5153 | 5325 | 11760 |
| White blood cell | WBC | 6721 | 2168 | 0 | 11055 |
| C-reactive Protein | CRP | 6212 | 182 | 2127 | 11766 |
| BMI | BMI | 6808 | 5131 | 5325 | 11744 |
| Diastolic bp | DBP | 6814 | 4610 | 0 | 11767 |
| PR interval | PR | 3433 | 745 | 0 | 11636 |
| QRS interval | QRS | 3433 | 752 | 0 | 11667 |
| eGFR | eGFR | 5627 | 0 | 0 | 11760 |
| HbA1c | HbA1c | 3 | 561 | 0 | 9515 |
| HDL | HDL | 6072 | 2117 | 2059 | 11767 |
| LDL | LDL | 5779 | 2042 | 2054 | 11561 |
| MCHC | MCHC | 1648 | 2168 | 0 | 11695 |
| Platelet count | PLT | 6706 | 2165 | 0 | 11686 |
| Systolic bp | SBP | 6819 | 4606 | 0 | 11772 |
| Total cholesterol | TC | 6070 | 2158 | 2072 | 11768 |
| Triglyceride | TG | 5857 | 2294 | 2066 | 11768 |
| CKD | CKD | 5627 (295) | 5154 (608) | 5325 (909) | 11768 (405) |
| Hypertension | Hypertension | 6716 | 5154 | 5325 | 11766 |
| Type II Diabetes | T2D | 6784 (2131) | 4566 (1780) | 4826 (2427) | 11388 (1702) |

**Table 2: Summary of Sample Sizes in Real Data by Phenotype and Cohort**

| **Method** | **N Causal Variants = 50** | **N Causal Variants = 200** |
| --- | --- | --- |
| GAUDI | 0.8591667 | 0.6729333 |
| HAUDI | 0.5522833 | 0.2953667 |
| LASSO | 0.5757667 | 0.3433333 |

**Table 3: Proportion of Estimated Effects Equal to Zero in Simulations**

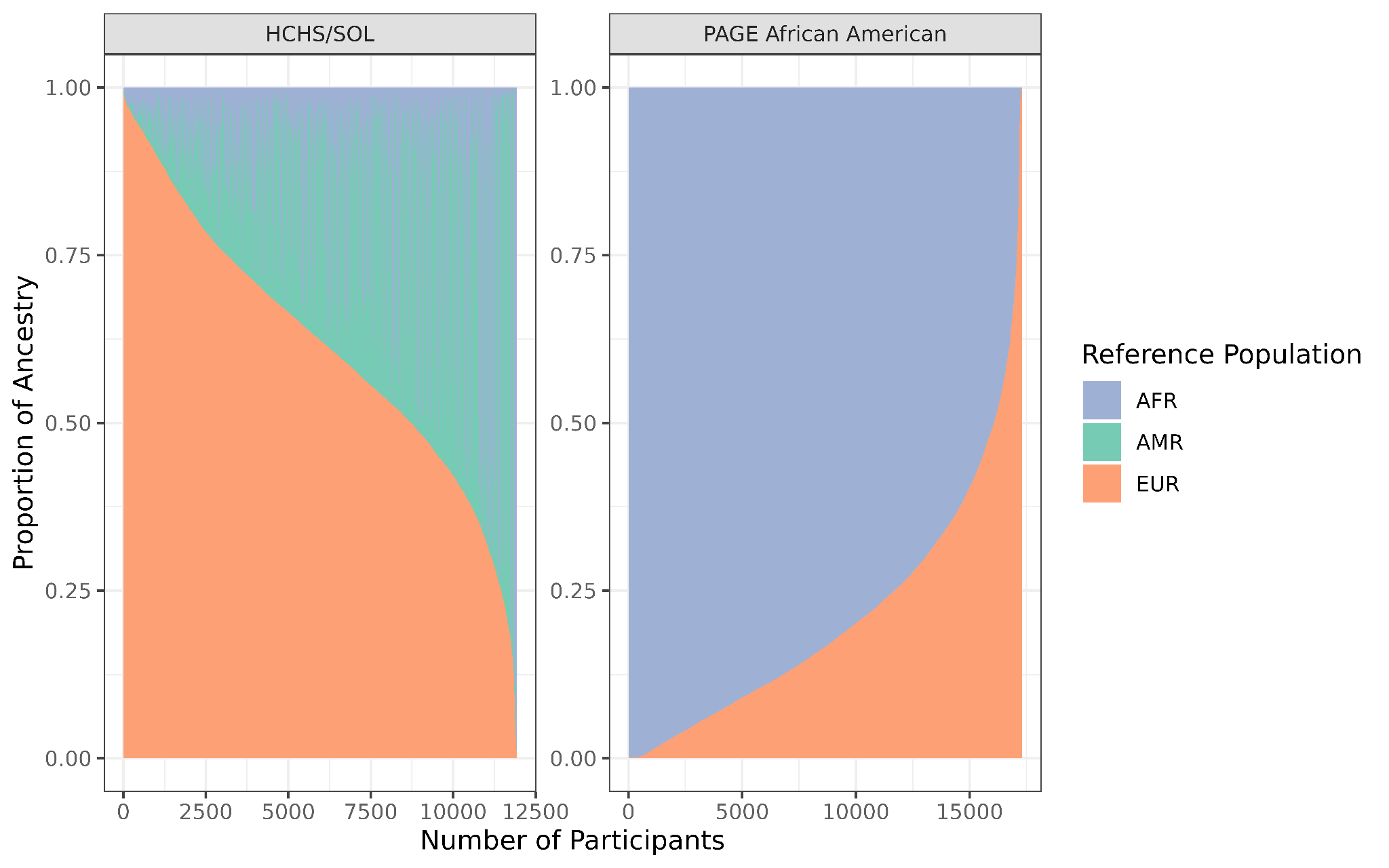

**Fig S1: Individual-Level Global Ancestry Proportions.** Average global ancestry proportions per-individual, estimated using FLARE.

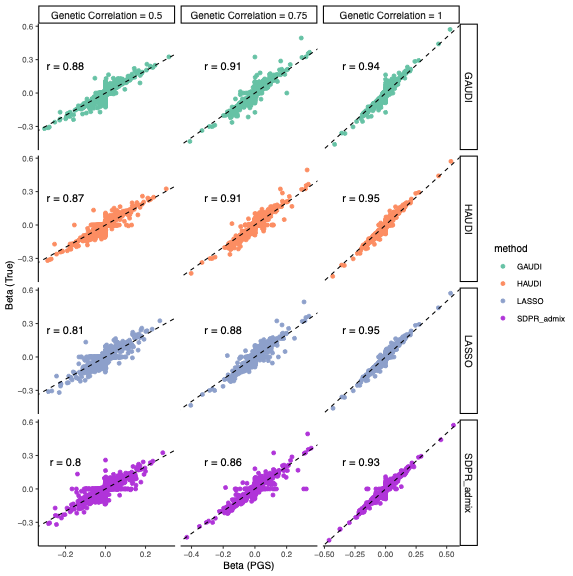

**Fig S2: Comparisons between estimated and true YRI-specific effects.** Plotted in each panel are the estimated (x axis) and true (y axis) variant effects, stratified by genetic correlation and PGS method. The correlation between the true and estimated effects (R) is annotated for each panel. This figure is restricted to YRI-specific effects. For LASSO, the x-axis shows estimated ancestry-agnostic effects

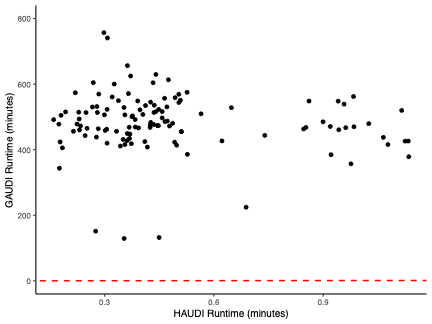

**Fig S3: Comparison between simulated HAUDI and GAUDI run time.** Plotted values correspond to the best-performing model for GAUDI and HAUDI for each combination of simulation settings. HAUDI and GAUDI runtimes are plotted on different scales due to large observed values for GAUDI. For purposes of comparison, we provide the dashed red line, with intercept of 0 and slope of 1.
